# Stress-Induced Cooperation Promotes Tolerance in Resource-Limited Auxotrophic Microbial Consortia

**DOI:** 10.64898/2026.01.08.698473

**Authors:** D Reyes-Gonzalez, H de Luna-Valenciano, F Santos-Escobar, M Rebolleda-Gomez, R Peña-Miller, A Fuentes-Hernandez

**Affiliations:** Programa de Biología Sintética, Centro de Ciencias Genómicas, Universidad Nacional Autónoma de México, 62210, Cuernavaca, México; Department of Ecology & Evolutionary Biology, University of California - Irvine, Irvine, California 92697; Programa de Biología de Sistemas, Centro de Ciencias Genómicas, Universidad Nacional Autónoma de México, 62210, Cuernavaca, México; Departamento de Biología, División de Ciencias Naturales y Exactas, Universidad de Guanajuato, 36050, Guanajuato, México

**Keywords:** Synthetic communities, Metabolic cooperation, Bacteriostatic antibiotics

## Abstract

Microbial communities often rely on metabolic cooperation, especially under nutrient limitation, but how antibiotics influence these interactions remains unclear. Here, we show that exposure to chloramphenicol, a ribosome-targeting bacteriostatic antibiotic, promotes cooperation in auxotrophic Escherichia coli consortia. Using a proteome-partition model, we find that chloramphenicol-induced growth inhibition elevates levels of the alarmone ppGpp, shifting proteome allocation toward amino acid biosynthesis and export. This regulatory response enhances cross-feeding and supports cooperative growth, even under antibiotic stress. Laboratory experiments confirm that, under low amino acid availability, co-cultures of leucine and phenylalanine auxotrophs display greater tolerance to chloramphenicol than monocultures. Both our theoretical and experimental results reveal a feedback loop in which antibiotic stress strengthens metabolic interdependence, buffering growth inhibition and enhancing community-level tolerance.

## 1 Introduction

Antibiotics are among the most powerful tools in medicine and microbiology, used primarily to treat infections by killing or inhibiting bacterial growth.^1^ However, in natural and clinical environments, bacteria rarely exist in isolation; rather, they inhabit complex communities structured by metabolic exchange, resource competition, and cooperative dependencies.^2, 3^ In these contexts, antibiotics can have broad and often unintended consequences. Beyond their direct action on individual cells, they can influence community structure by altering growth rates and shifting the balance between competitive and cooperative interactions.^4, 5^ At subinhibitory concentrations, antibiotics have been shown to modify gene expression and metabolism across species.^6, 7^ These changes can weaken or reinforce interdependencies and ultimately affect the stability and evolutionary dynamics of microbial communities.^8^

Importantly, the structure of microbial communities can also shape the outcomes of antibiotic exposure. Resistant strains may protect susceptible partners by degrading or sequestering antibiotics,^9^ while other interspecies interactions can destabilize communities by shifting selective pressures.^5^ For instance, cooperative interactions can be disrupted under stress, as mutualistic dependencies become liabilities when partners are lost.^10^ Ecological theory predicts that mutualistic consortia may be particularly susceptible to antibiotics due to their metabolic interdependence, while competitive interactions can facilitate competitive release and promote the expansion of resistant strains.^4, 11, 12^ Ecological context, therefore, plays a central role in shaping the efficacy of antibiotic treatment, influencing not only susceptibility but also how bacterial interactions modulate community responses.^13^

Among the ecological interactions that influence microbial responses to antibiotics, syntrophic cooperation among auxotrophic strains is a well-characterized and mechanistically tractable model system.^14, 15^ Auxotrophy, the inability to synthesize specific essential metabolites such as amino acids^16^ and vitamins,^17^ can lead to mutualistic consortia in which strains exchange metabolites to sustain growth. These metabolic interdependencies are widespread in both natural microbiomes^18, 19^ and engineered systems,^18–21^ where they support division of labor strategies^22^ and promote resilience under environmental stress.^23–25^ Under such conditions, growth-arrested cells can release shared metabolites that sustain surrounding populations, reinforcing community stability and survival.^26^ Metabolite exchange can also modulate antibiotic susceptibility, and in some cases, enable collective tolerance in otherwise sensitive populations.^27–29^

While cross-feeding is known to influence antibiotic susceptibility, it remains unclear how antibiotics themselves reshape cooperative interactions in auxotrophic consortia. Here, we focus on chloramphenicol (CHL), a ribosome-targeting bacteriostatic antibiotic, to ask whether sublethal translational inhibition can enhance cooperation in metabolically interdependent bacteria. We hypothesized that CHL promotes metabolic cooperation by elevating ppGpp, the alarmone guanosine tetraphosphate that coordinates the stringent response.^30, 31^ This shift in proteome allocation enhances amino acid biosynthesis and export, reinforcing interdependence and enabling collective growth under antibiotic stress. To test this, we combined theoretical modeling with co-culture experiments using auxotrophic *Escherichia coli* K12 exposed to gradients of amino acids and CHL. Our results show that under nutrient limitation, CHL amplifies cross-feeding and sustains co-culture growth, yielding greater tolerance than monocultures. This emergent tolerance reflects regulatory feedbacks in which antibiotic-induced growth inhibition reinforces cooperative metabolism and promotes community-level resilience.

## 2 Results

Understanding how CHL modifies cooperation in auxotrophic bacteria requires a framework that links intracellular physiology to population dynamics. Proteome partition models provide such a framework by describing how cells distribute limited biosynthetic capacity among metabolism and stress-related functions under different environmental conditions.^32, 33^ These models capture regulatory constraints that operate across timescales and can be extended to explore how intracellular resource allocation influences ecological interactions.^34–36^ In earlier work,^37^ we developed a multiscale model that connects proteome regulation to growth and amino acid exchange in auxotrophic *E. coli*. The model links ppGpp-controlled proteome allocation to the production and release of essential amino acids, predicting when and how metabolic cooperation emerges under nutrient limitation. Here, we apply this framework to evaluate how a bacteriostatic antibiotic that inhibits translation by blocking peptidyl-transferase activity in the 50S ribosomal subunit^38^ affects proteome allocation and modulates metabolic interactions.

Following the formulation of Scott et al.,^39^ the proteome is represented as three sectors. The ribosomal sector (R) determines translational capacity, the enzymatic sector (E) includes metabolic and biosynthetic proteins, and the growth-independent sector (Q) captures housekeeping and stress-related functions. To explicitly model auxotrophy, the enzymatic sector is subdivided into the amino acid biosynthesis fraction (*E*_aa_) and all other metabolic functions, such that the total proteome satisfies

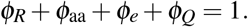

Proteome allocation is controlled by ppGpp, which accumulates when ribosomes stall due to nutrient limitation^40^ or CHL binding.^41^ Elevated ppGpp reduces ribosome synthesis and increases transcription of biosynthetic enzymes, thereby reallocating proteome investment away from translation and toward metabolism, including amino acid biosynthesis. This process links translational inhibition to metabolic output and, consequently, to cooperative interactions in auxotrophic consortia. CHL is incorporated by distinguishing active ribosomes from drug-bound inhibited ribosomes, allowing ribosomal inhibition to reduce effective translation and increase ppGpp levels. A description of the model formulation is provided in the Methods.

### 2.1 ppGpp Regulation by Glucose and Amino Acid Levels

We used numerical simulations to explore how the proteome allocation model responds to gradients of glucose, external amino acids, and CHL, based on the parameters in Table S1. For each environmental combination, the numerical experiment produced a proteome allocation profile from which we computed the growth rate (*λ*) and amino acid export rate (*β*), as illustrated in Figure 1A. Under low amino acid availability (Figure 1B), ribosome stalling increased ppGpp, which reduced ribosomal mRNA (*m*_*R*_) and enhanced metabolic transcription (*m*_*E*_). As a result, the proteome shifted from the ribosomal sector (*φ*_*R*_) toward metabolism (*φ*_*E*_), with an increase in the amino acid biosynthesis sub-sector 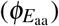. This redistribution increased amino acid export rate (*β*) because cells invested more in internal amino acid biosynthesis. In contrast, when amino acids were externally supplied (Figure 1C), ppGpp levels remained low, the proteome shifted toward ribosome synthesis, and growth rate increased. Under high-amino acid conditions, amino acid export rates declined because biosynthetic investment was reduced.

**Figure 1.**
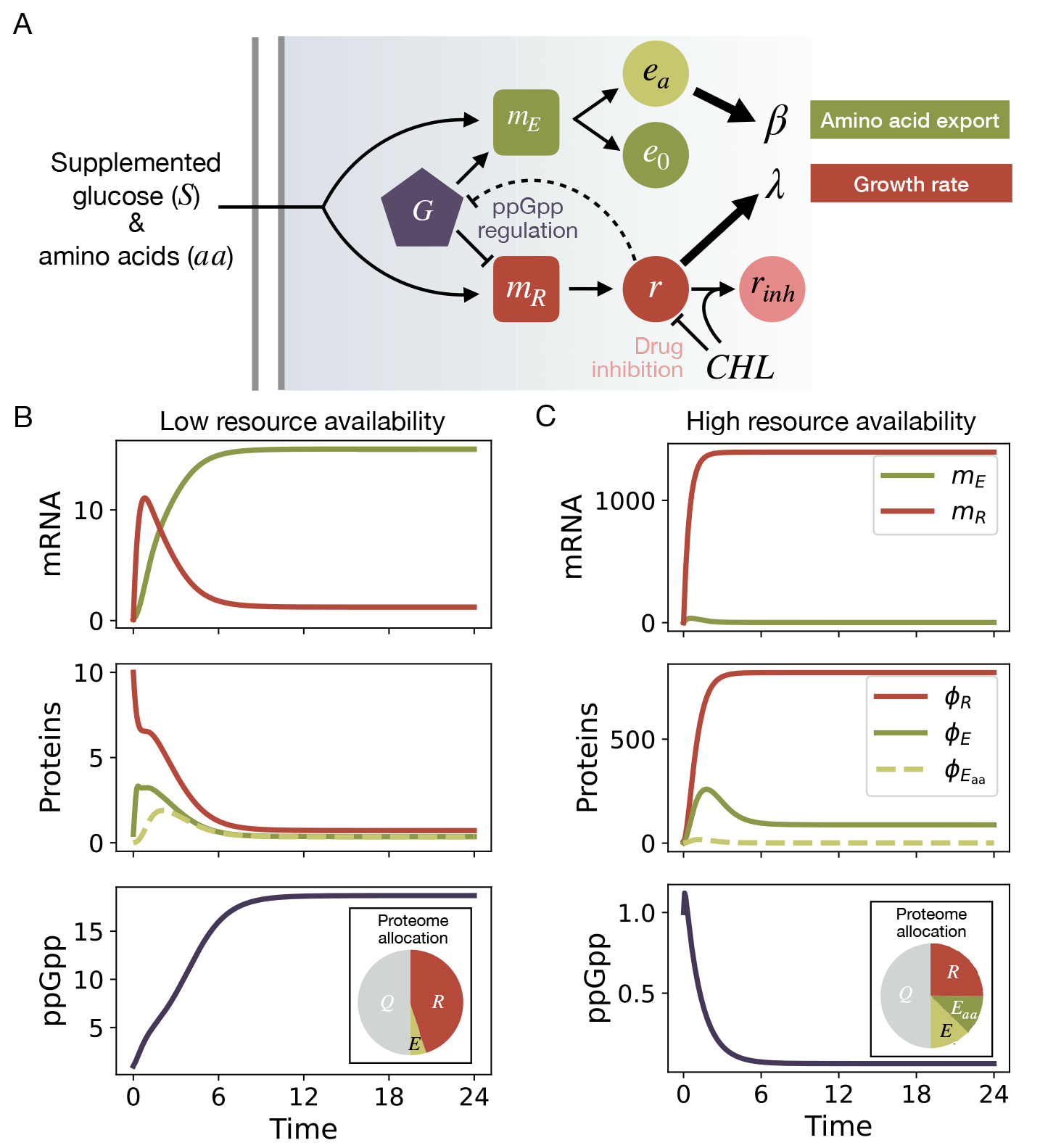
Proteome partition model. (A) Diagram of the proteome allocation model. The model describes how nutrient availability and *CHL* affect cellular physiology via ppGpp-mediated regulation. Squares represent mRNAs (*m*_*E*_, *m*_*R*_), circles denote proteins (*e*_*o*_, *e*_*a*_, *r, r*_inh_), and the pentagon represents ppGpp (*g*). Nutrients (*S, aa*) activate transcription; ppGpp promotes *m*_*E*_ and represses *m*_*R*_. Translation yields metabolic enzymes (*e*_*o*_, *e*_*a*_) and ribosomes (*r*), which are inhibited by *CHL* to form *r*_inh_. Ribosome abundance regulates ppGpp synthesis. Growth rate (*λ*) and cooperative output (*β*) are proportional to *r* and *e*_*a*_, respectively. Pointed arrows indicate activation or production; blunt arrows indicate inhibition. (B) Simulated dynamics of mRNA expression, proteome allocation, and ppGpp concentration under low-resource conditions (*S* = 1.0, *aa* = 0.1). Top panel: mRNA concentrations of the metabolic (*m*_*E*_) and ribosomal (*m*_*R*_) sectors over time. Middle panel: proteome fractions allocated to ribosomes (*φ*_*R*_), total metabolic enzymes (*φ*_*E*_), and amino acid biosynthesis 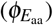. Bottom panel: intracellular concentration of the alarmone ppGpp (*g*), which increases as growth slows. (C) Simulated dynamics under high-resource conditions (*S* = 1.0, *aa* = 10.0). Top panel: elevated amino acid availability suppresses ppGpp-regulated expression of metabolic genes, reducing *m*_*E*_ while enhancing *m*_*R*_. Middle panel: proteome shifts toward ribosome synthesis (*φ*_*R*_), with reduced allocation to amino acid biosynthesis 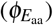. Bottom panel: ppGpp concentration (*g*) remains low, consistent with faster growth. Inset: pie chart showing proteome allocation into ribosomal (*R*), enzymatic (*E* and *E*_*aa*_), and housekeeping (*Q*) sectors as obtained from model simulations.

A similar regulatory pattern emerged along glucose gradients. Increasing glucose availability elevated ribosomal mRNA production and expanded the ribosomal proteome fraction (Figure S1). ppGpp levels remained low, supporting rapid translation and fast growth but limiting the proteome available for biosynthetic enzymes. Thus, carbon-rich conditions favored autonomous growth rather than cooperation. Supplementing external amino acids reproduced a transition in regulatory state (Figure S2): as external amino acid concentrations increased, ppGpp levels declined, shifting proteome allocation toward ribosome synthesis, increasing growth rate (*λ*), and reducing amino acid export (*β*).

In summary, the model explores how ppGpp regulation could integrate signals from nutrient limitation and translational stress to shape proteome allocation. It predicts that transcription shifts away from ribosomal genes and toward biosynthesis and stress response. As a result, cooperative output is modulated by environmental conditions, linking intracellular regulation to metabolite exchange. Specifically, modeling results suggest that moderate stress or nutrient limitation may enhance metabolite production and cooperation, whereas abundant resources could suppress amino acid export and reduce cross-feeding.

### 2.2 Chloramphenicol Effects on Growth and Amino Acid Export

We next asked how translational inhibition by CHL interacts with nutrient availability to reshape the trade-off between growth and amino acid export rates. Model simulations revealed that ppGpp dynamics depend strongly on external amino acid availability: at high supplementation, ppGpp remained suppressed, while at lower concentrations, trajectories crossed a threshold and converged to a high-ppGpp state, indicating bistability in response to nutrient stress (Figure 2A). This regulatory shift suggests that under nutrient-rich conditions, faster translation may render cells more vulnerable to CHL due to increased sensitivity to ribosomal inhibition.^42^

**Figure 2.**
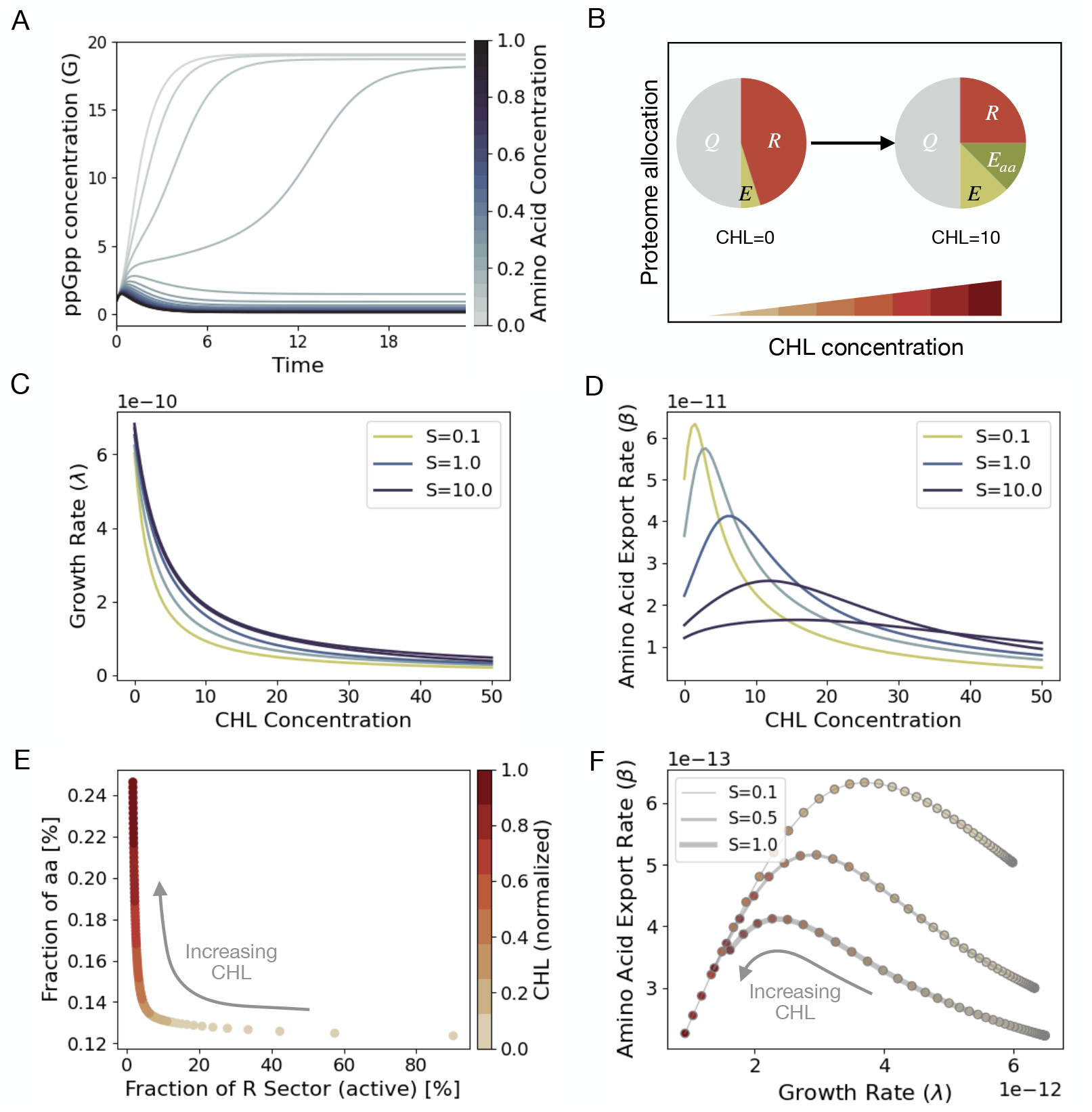
Modeling the effects of CHL on ppGpp dynamics and proteome allocation. (A) Time courses of ppGpp (*g*) for increasing CHL concentrations. Higher CHL slows translation and leads to elevated ppGpp, reflecting stronger activation of the stringent response. (B) Proteome composition at the end of the simulation without CHL (left) and with 10 µg/mL CHL (right). CHL reduces the ribosomal proteome fraction (*φ*_*R*_) and increases allocation to amino acid biosynthesis 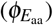, resulting in lower growth rate (*λ*) and higher amino acid export (*β*). (C) Growth rate *λ* across CHL concentrations for a range of substrate levels (*S* = 0.1–10, log-spaced), with amino acids held constant (*aa* = 1). Higher *S* increases growth potential but preserves the decline in *λ* with CHL. (D) Amino acid export rate *β* across the same CHL and *S* gradients as in panel (C). CHL produces an increase in *β* at intermediate doses, followed by a decline at higher concentrations, and the magnitude of this response depends on carbon availability. (E) Growth rate (*λ*, left) and amino acid export rate (*β*, right) across a CHL gradient for different substrate concentrations (*S*). At low CHL, growth dominates; at intermediate CHL, ppGpp-driven proteome reallocation increases *β*; at high CHL, both *λ* and *β* decline due to strong translational inhibition. These effects are most pronounced under low-resource conditions. (F) Joint relationship between *λ* and *β* across CHL concentrations (colour gradient) for different *S* levels (line thickness). Increasing CHL shifts cells along a trade-off trajectory in which proteome resources move from ribosomal activity toward amino acid biosynthesis. The trade-off is strongest at low *S*, where biosynthetic investment increases in response to CHL.

Increasing CHL expands the pool of inactive ribosomes (*r*_inh_) while elevating ppGpp, which suppresses ribosome synthesis and redirects proteome capacity toward metabolism (Figure 2B). This reallocation increases the amino acid biosynthesis sub-sector 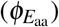, enhancing export rates (*β*) at low CHL concentrations. At higher CHL doses, however, ribosome inhibition becomes the main constraint, broadly impairing biosynthesis and suppressing both growth and metabolite export (Figure S3) I.

To examine this trade-off in more detail, we compared the trajectories of growth and export across a range of CHL concentrations and nutrient levels. Figure 2C–D illustrates how *λ* decreases monotonically with CHL concentration, reflecting reduced growth across carbon gradients (*S* = 0.1 to 10), while *β* shows a non-monotonic pattern, with export rates declining at high CHL concentrations but increasing at intermediate levels. Plotting the proteome allocation across CHL concentrations reveals how increasing drug levels reduce the ribosomal fraction while expanding the amino acid biosynthesis sector (Figure 2E). This trade-off results in a non-monotonic trajectory of cooperative output: while growth rate (*λ*) declines steadily, amino acid export (*β*) rises to a maximum before collapsing at high CHL concentrations. The magnitude of this effect depends on nutrient availability, with the strongest cooperative enhancement occurring under carbon limitation (Figure 2F). In summary, the model predicts that bacteriostatic inhibition of translation reconfigures intracellular regulation to transiently support cooperative metabolism, suggesting that physiological constraints at the cellular level can generate emergent community-level behaviors.

### 2.4 Experimental System: Auxotrophic Microbial Consortia

Our model suggests that under translational stress, proteome reallocation toward amino acid biosynthesis can enhance cooperative output. To test this prediction experimentally, we used a synthetic consortium of two *E. coli* auxotrophs from the Keio collection:^43^ Δ*leuB* (leucine auxotroph) and Δ*pheA* (phenylalanine auxotroph). We fluorescently labeled Δ*pheA* with mCherry and Δ*leuB* with eCFP, enabling quantification of their relative abundances by flow cytometry. These strains obligatorily cross-feed the amino acids they cannot synthesize:^37^ Δ*pheA* secretes leucine consumed by Δ*leuB*, while Δ*leuB* secretes phenylalanine required by Δ*pheA*, with glucose as the sole carbon source (Figure 3A).

**Figure 3.**
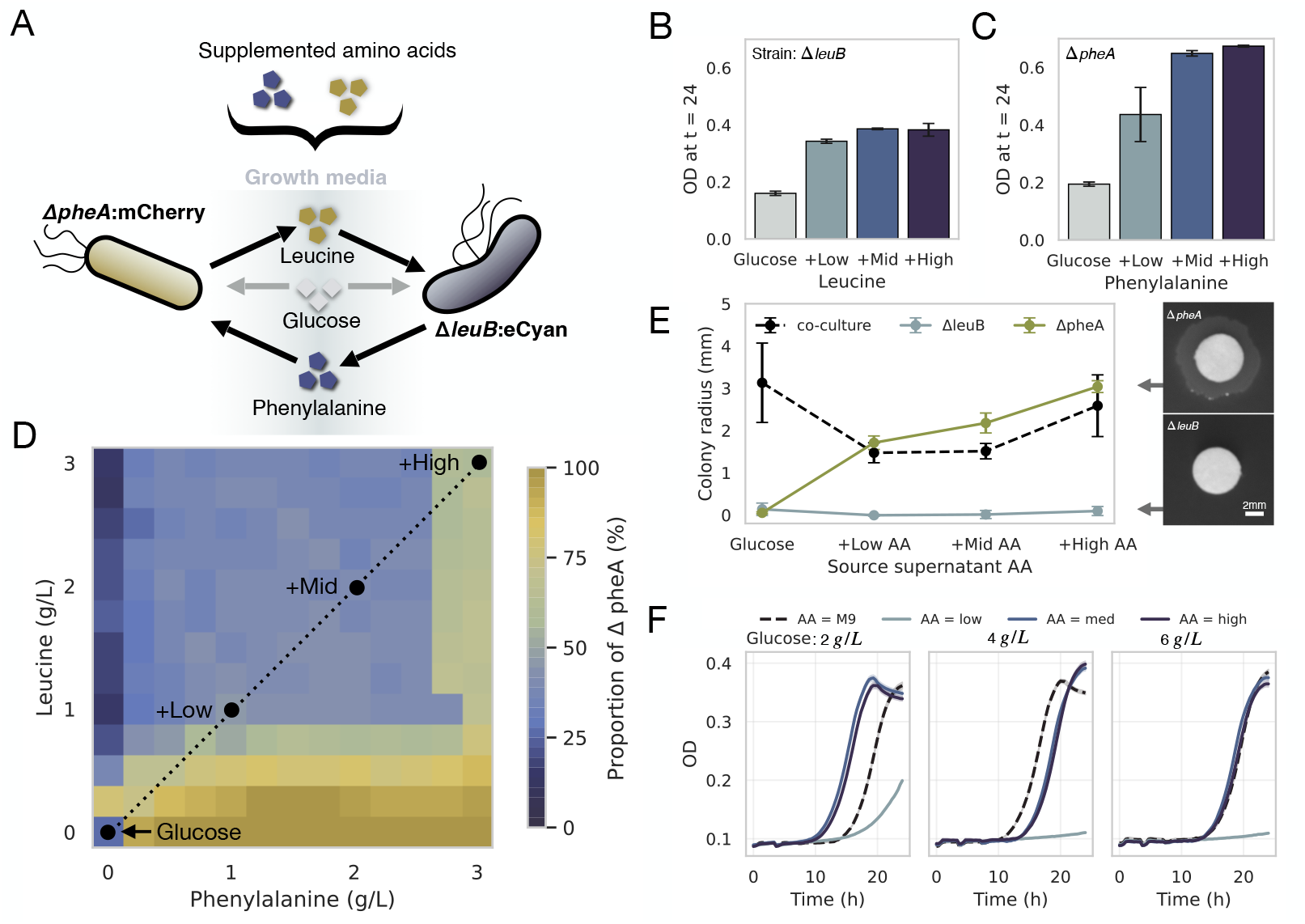
Experimental model system. (A) Schematic of the co-culture system. Auxotrophic *E. coli* strains Δ*pheA* (yellow) and Δ*leuB* (blue) exchange amino acids: Δ*pheA* exports leucine consumed by Δ*leuB*, while Δ*leuB* produces phenylalanine consumed by Δ*pheA*. Both strains use glucose as the sole carbon source. (B) Final densities of Δ*pheA* in monoculture across a range of supplemented phenylalanine concentrations, measured as optical density (OD_630_) at 24 h. Colors indicate increasing supplementation: no amino acid (gray), low to high (light to dark blue). Bars show mean ± SEM. (C) Final densities of Δ*leuB* in monoculture across leucine supplementation levels, colored as in (B). (D) Checkerboard co-culture competition assay. Heatmap shows the final proportion of Δ*pheA* (%) after 48 h, starting from equal initial frequencies, as a function of supplemented phenylalanine (x-axis) and leucine (y-axis). Strain frequencies were quantified by flow cytometry using red and cyan fluorescence. Dotted lines indicate the amino acid combinations (none, low, mid, high) used in subsequent experiments. (E) Colony radius on M9 agar for lawns of co-culture (black dashed), Δ*leuB* (blue), and Δ*pheA* (yellow), after applying supernatants from liquid cultures grown in M9, Low, Mid, or High amino acid supplementation. (F) Growth curves of the Δ*pheA*–Δ*leuB* co-culture under varying amino acid and glucose concentrations. Black dotted line shows growth in M9 media; supplemented amino acid conditions are indicated with a blue gradient. Each panel shows a different glucose concentration: from left to right, 2, 4, and 6 g/L.

To establish this auxotrophic pair as a tunable system for metabolic cooperation,^15^ we first confirmed their individual amino acid requirements. In monoculture without supplementation, both Δ*pheA* and Δ*leuB* exhibited minimal growth. Adding the respective amino acid increased optical density, followed a saturating dose–response relationship (Figures 3B–C). These results confirm that growth is limited by amino acid availability and validate the auxotrophic dependencies. We therefore defined a supplementation gradient consisting of Glucose (M9 + glucose only), +Low (basal), +Mid (2 *×* basal), and +High (3 *×* basal), based on these dose–response curves and predictions from a genome-scale model.^37^ Detailed concentrations and calculations are provided in the Methods.

Next, we examined co-culture dynamics across a matrix of supplemented amino acid concentrations (Figure 3D). When leucine was omitted, the population was composed primarily of Δ*pheA* (mean frequency: 93.1 *±* 20.2%, SD, n = 13), whereas phenylalanine omission resulted in a higher frequency of Δ*leuB* (Δ*pheA* frequency: 25.0 *±* 17.2%, SD, n = 13), confirming the auxotrophic dependencies of each strain. At low supplementation of both amino acids, the two strains maintained comparable frequencies (Δ*pheA*: 67.7 *±* 29.1%), consistent with mutual dependence and bidirectional crossfeeding. As external amino acid availability increased to Mid and High levels, the frequency of Δ*pheA* progressively declined (45.8% and 39.4%, respectively), indicating a shift in population structure driven by reduced biosynthetic constraints. This pattern reflects a transition from mutualistic to competitive dynamics as the need for cross-feeding is alleviated by external supplementation.^21–23^

To verify that cooperative growth is mediated by metabolic exchange, we tested whether spent media from co-cultures could support growth in the absence of supplemented amino acids. Cell-free supernatants from 24-hour cultures were applied to filter disks on M9-glucose agar without amino acid supplementation (Figure 3E). Supernatants from Δ*pheA* or Δ*leuB* monocultures failed to support co-culture growth, confirming that each strain releases only one of the required amino acids and that both are needed to sustain the consortium. In contrast, supernatant from the unsupplemented co-culture supported robust colony expansion (mean colony radius: 3.13 *±* 0.94 mm), consistent with the accumulation of both amino acids during liquid growth. When monocultures were supplemented with increasing concentrations of both amino acids, only the Δ*pheA* supernatant supported growth (1.71 *±* 0.16 mm at Low AA, 3.04 *±* 0.14 mm at High AA), whereas Δ*leuB* supernatants did not (≤ 0.09 *±* 0.11 mm across all conditions).

Together, these results show that the auxotrophic consortium exhibits nutrient-dependent interaction dynamics: under amino acid limitation, reciprocal cooperation supports coexistence, while high-resource environments and disrupt metabolic interdependence. This tunable experimental system, in which amino acid and carbon availability modulate the balance between cooperation and competition, provides a controlled framework to examine how chloramphenicol influences metabolic interactions and drug susceptibility profiles.

### 2.3 Dynamic Interaction Profiles Driven by Resource Availability

To examine how nutrient availability shapes ecological interactions in the auxotrophic consortium, we quantified strain frequencies and total biomass after 48 h of co-culture across a matrix of leucine and phenylalanine concentrations. At low amino acid supplementation, both strains remained at similar frequencies, consistent with mutual dependence and reciprocal cross-feeding (Figure 3D). As external amino acid supply increased, this dependence weakened, and strain composition progressively shifted toward Δ*leuB*, reflecting a transition from mutualism to competitive exclusion.

We next asked how carbon availability interacts with amino acid supplementation to modulate cooperative dynamics. Co-cultures were grown across a gradient of glucose concentrations under four defined amino acid regimes: M9 (no supplementation), Low, Mid, and High. At high amino acid supplementation, co-cultures achieved high final densities, which further increased with rising glucose concentrations. In contrast, under Low supplementation, co-cultures exhibited reduced growth and reached low final densities. Increasing glucose under these conditions further decreased final density, suggesting that limited amino acid availability combined with excess carbon destabilizes cooperation and leads to population collapse.

Strikingly, in the complete absence of amino acid supplementation, co-cultures still reached substantial densities, although with a prolonged lag phase compared to Mid and High conditions (Welch’s t-test; H_0_: equal lag time; p < 0.01). As glucose concentrations increased, growth rate in the unsupplemented condition also increased and eventually approached the levels observed under Mid and High supplementation. At 4 g/L glucose, the growth rate in M9 was significantly higher than in both Mid and High supplementation conditions (Welch’s t-test; p < 0.001), indicating that cooperative metabolism can outperform externally relieved biosynthetic constraints under intermediate carbon availability. At 6 g/L glucose, growth rates across all supplementation regimes were statistically indistinguishable (Welch’s t-test; H_0_: equal means; p > 0.05), consistent with the interpretation that biosynthetic constraints are fully relieved at high resource availability and cooperative exchange becomes dispensable.

### 2.4 Ribosome Inhibition Promotes Cooperation in Auxotrophic Consortia

We next investigated how ribosome inhibition alters interaction dynamics within the auxotrophic consortium. Monocultures and co-cultures of Δ*leuB* and Δ*pheA* were exposed to a range of CHL concentrations (0–15 *µ*g/mL) in M9 minimal medium with glucose for 48 h. Cultures were grown in 96-well plates and monitored by plate reader, which recorded OD and fluorescence every 20 minutes. Final co-culture composition was quantified by flow cytometry (Figure 4A), allowing us to assess both growth and strain frequencies across the CHL gradient and amino acid supplementation levels.

**Figure 4.**
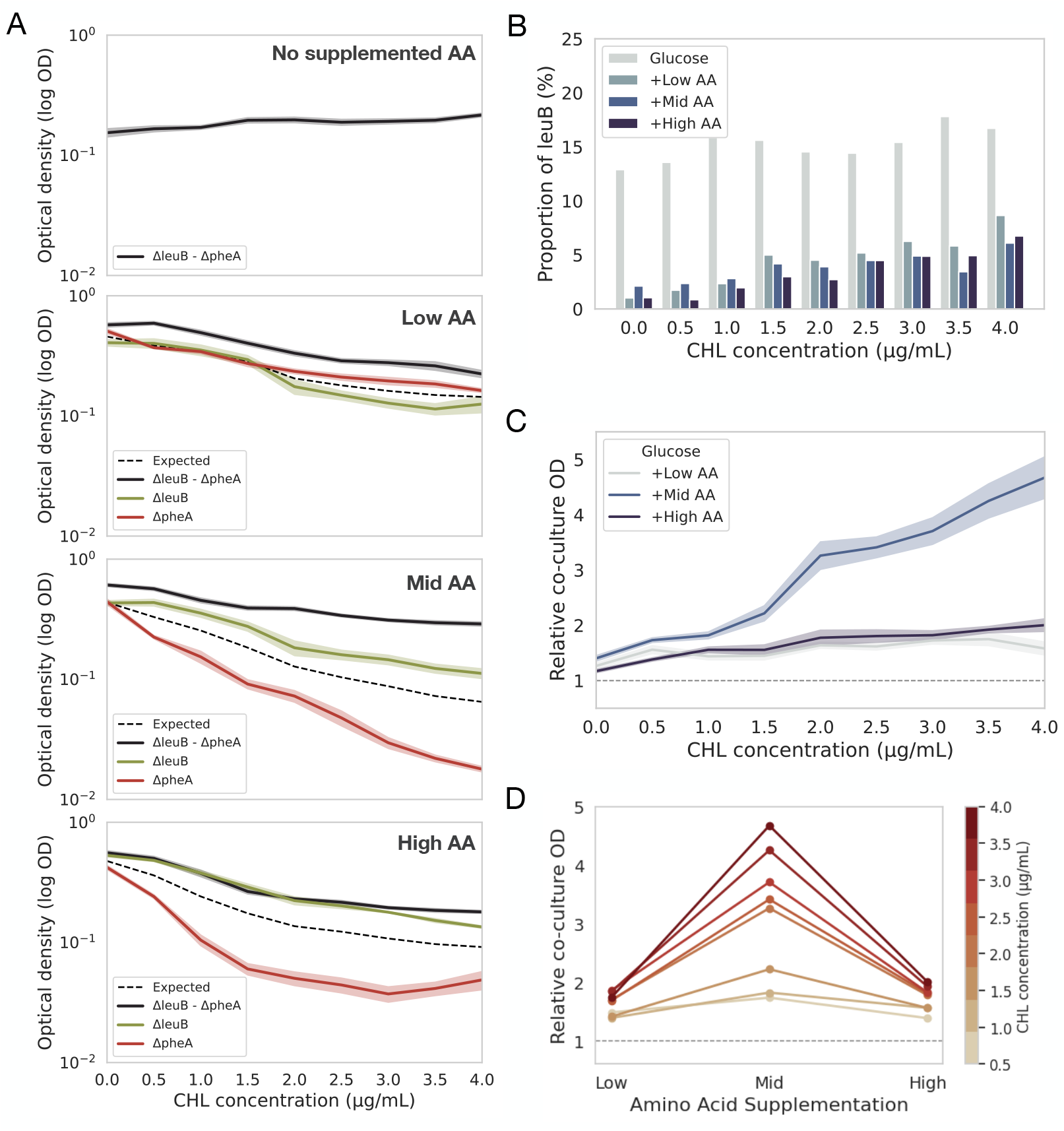
Growth Inhibition by Chloramphenicol Enhances Cooperation in Auxotrophic Co-cultures. (A) Optical density (OD) after 48h growth in M9-glucose medium with increasing concentrations of CHL for Δ*leuB* (green), Δ*pheA* (red), and the co-culture Δ*leuB*-Δ*pheA* (solid black). Thick lines denote the mean and areas the standard deviation. The dashed black line indicates the expected OD under an additive model (mean of monocultures). (B) Fraction of the eCyan-labeled strain (Δ*leuB*) in co-culture with Δ*pheA* measured using flow cytometry across a range of CHL concentrations. Each bar corresponds to distinct amino acid supplementation levels (No supplementation, Low, Mid, and High). (C) Fold change in final OD of the Δ*leuB*-Δ*pheA* co-culture relative to the expected OD (mean of monocultures) at increasing CHL concentrations. Each thin line corresponds to a biological replicate; the thick black line indicates the mean. The dashed gray line at fold change=1 marks the additive expectation. Values above 1 indicate increased yield in co-culture compared to monocultures. (D) Fold change in OD of the Δ*leuB*-Δ*pheA* co-culture relative to the expected monoculture average across amino acid supplementation levels (Low, Mid, High) and CHL concentrations (0.5–15*µ*g/mL). Co-cultures consistently outperformed monocultures, with the cooperative benefit increasing with CHL and reaching a maximum at intermediate amino acid levels.

Under Low amino acid supplementation, strain frequencies remained balanced across CHL concentrations, consistent with comparable susceptibility of Δ*leuB* and Δ*pheA*. At Mid and High supplementation, CHL exposure shifted the relative abundance toward Δ*leuB*, indicating that resistance differences emerge as amino acid availability increases (Figure 4B). This pattern is supported by IC50 estimates: susceptibility did not differ at Low supplementation (Mann Whitney test, *p* = 0.72), whereas at higher supplementation the IC50 values diverged, with a statistically significant difference at High AA (Mann Whitney test, *p <* 0.001; Figure S4).

We next compared co-culture growth to the additive expectation obtained by averaging the two monocultures under each amino acid condition and CHL concentration (Figure 4C). Co-cultures consistently exceeded this non-interaction benchmark, and the magnitude of the deviation depended on nutrient supply, with the largest effect at intermediate supplementation. Across all amino acid conditions, relative co-culture density increased with CHL concentration (Figure 4D), consistent with model predictions that translational inhibition under nutrient limitation elevates ppGpp, redirects proteome allocation toward biosynthesis, and increases amino acid export (Figure 2C).

To directly assess whether CHL modulates cooperative capacity, we tested whether supernatants from treated cultures could support community growth on minimal media. Cell-free supernatants from 24-hour cultures were applied to filter disks on M9-glucose agar without amino acid supplements (Figure S5). In monocultures grown with amino acid supplementation, increasing CHL concentrations progressively reduced colony radii, indicating a dose-dependent decline in their capacity to support co-culture expansion. For instance, Δ*pheA* supernatants under High supplementation decreased from 3.04 mm to 1.94 mm as CHL increased from 0 to 3 *µ*g/mL. Supernatants from Δ*leuB* remained consistently low across all CHL concentrations (≤ 0.13 mm). In co-cultures grown without amino acid supplementation, supernatants supported colony expansion in the absence of CHL (3.13 mm), indicating accumulation of diffusible metabolites. Addition of 0.5 *µ*g/mL CHL reduced colony size to 2.07 mm. At higher CHL concentrations, colony size increased, reaching 2.28 mm at 1.5 *µ*g/mL and 3.27 mm at 3 *µ*g/mL, consistent with enhanced metabolite release under translational inhibition. (Figure 2F).

### 2.5 Cooperation Enhances CHL Tolerance Under Amino Acid Limitation

To determine how nutrient conditions influence CHL tolerance, we measured the optical density of the Δ*leuB*-Δ*pheA* co-culture after 48 hours across a matrix of amino acid supplementation levels and CHL concentrations (Figure 5A). At each CHL dose, we asked which amino acid concentration supported the highest final density. Across the entire CHL gradient, including in the absence of the drug, intermediate supplementation (Mid) consistently supported the greatest co-culture growth. This nonlinear growth pattern was specific to co-cultures. In monocultures, Δ*pheA* showed high CHL tolerance under Low supplementation but became increasingly susceptible as amino acid availability increased. Δ*leuB* exhibited relatively stable sensitivity across nutrient conditions. These contrasting profiles confirm that the co-culture phenotype is not a simple average of monoculture responses but emerges from nutrient-dependent metabolic interdependence.

**Figure 5.**
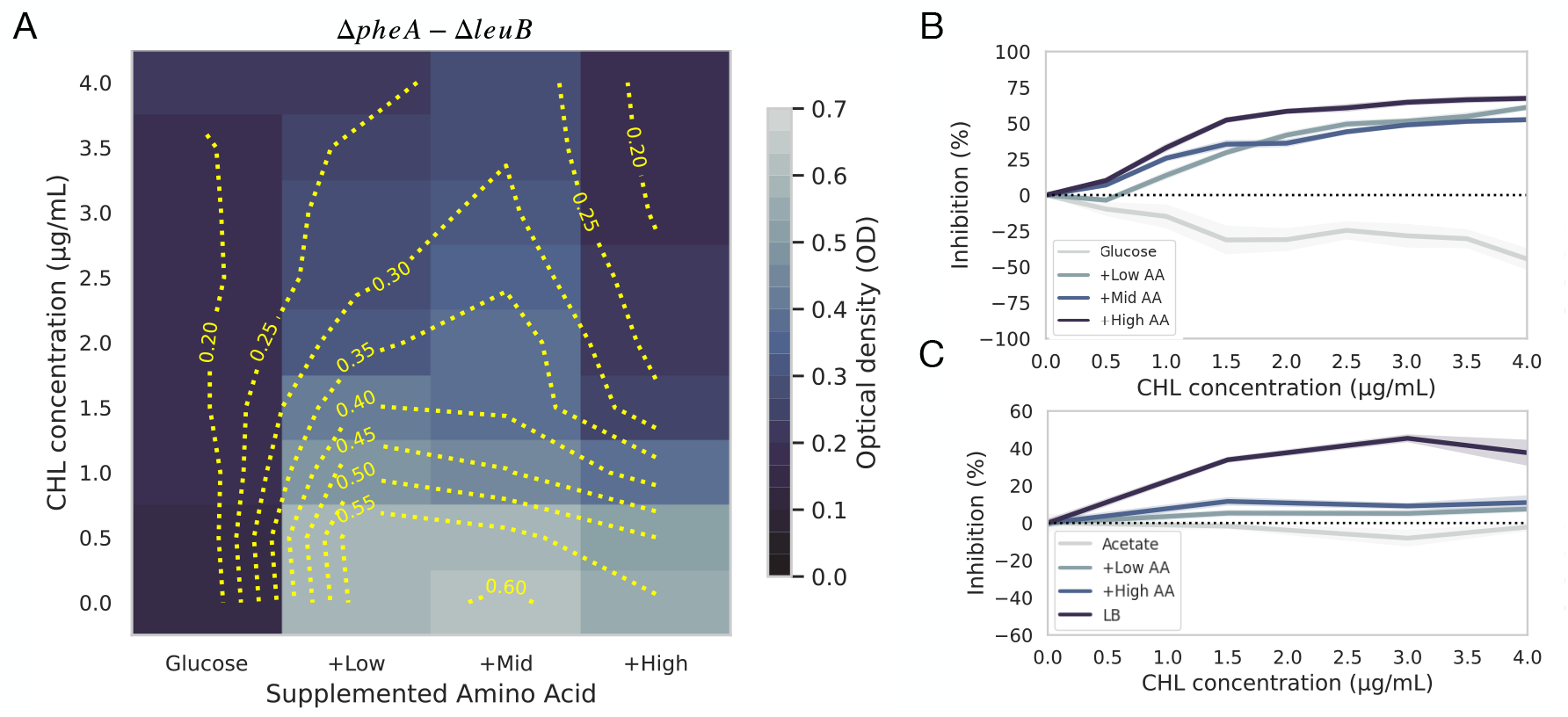
CHL tolerance is enhanced by metabolic cooperation under nutrient limitation. (A) Optical density (OD) of the Δ*leuB*–Δ*pheA* co-culture after 48 h across a matrix of amino acid supplementation levels (x-axis) and CHL concentrations (y-axis). Values are shown as a heatmap, and yellow contour lines mark fixed OD levels (0.10–0.60) to emphasize non-linear growth responses. Under low amino acid supply, OD increases at intermediate CHL concentrations relative to the CHL-free condition, indicating enhanced growth under moderate translational inhibition. At higher amino acid supplementation, CHL produces a monotonic decline in OD, consistent with conditions where cooperation is not required and growth is limited primarily by ribosomal inhibition. (B) CHL inhibition curves for the Δ*leuB*–Δ*pheA* co-culture across four amino acid supplementation regimes. Percent inhibition was calculated relative to the no-drug control (0 *µ*g/mL CHL) for each amino acid background. Curves show the mean inhibition response, with shaded bands indicating the SEM across biological replicates. Nutrient conditions are shown using a fixed color scheme: grey for glucose alone, and a graded blue palette for low, mid, and high amino acid supplementation. Without amino acid supplementation, CHL produces negative inhibition that becomes more pronounced as the drug concentration increases. In contrast, amino acid supplementation restores the expected monotonic increase in inhibition with rising CHL concentrations. (C) CHL inhibition curves for the co-culture in acetate-based medium. Lines show responses in acetate alone (grey), acetate + low amino acids (light blue), acetate + high amino acids (mid blue), and LB (dark blue). Inhibition values are plotted relative to the no-drug control for each condition, and shaded areas indicate the variability across replicates.

Drug-response curves summarized these trends (Figure 5B). Inhibition was calculated as the percent reduction in final optical density relative to the no-drug control, such that values above zero indicate growth inhibition and values below zero indicate enhanced growth. In glucose-only media without amino acid supplementation, CHL produced negative inhibition values that became more pronounced with increasing drug concentration, indicating that the co-culture grew better with CHL than without it. Adding amino acids eliminated this effect and restored a monotonic inhibition profile.

We next examined whether this cooperative response depends on the carbon source (Figure 5C). In acetate-based media, which supports slow growth, the co-culture again exhibited negative inhibition in the absence of amino acids. Supplementing acetate with amino acids restored a monotonic inhibition pattern. In nutrient-rich LB, populations were strongly inhibited at all CHL concentrations. These results suggest that enhanced growth occurs specifically under nutrient-limited or slow-growth regimes, conditions the model predicts elevate ppGpp levels and increase biosynthetic allocation.

Together, these findings show that CHL promotes co-culture growth only when metabolic interdependence is required. Under nutrient limitation, moderate translational inhibition improves growth, producing negative inhibition values in minimal media. When amino acid supplementation relieves biosynthetic constraints, CHL leads to a typical dose-dependent inhibition pattern. These observations suggest that antibiotic tolerance can arise from a physiological shift in resource allocation that strengthens cooperative metabolism, allowing mixed populations to better endure translational stress than monocultures.

## Discussion

Microbial communities are structured by a complex network of interactions that arise from metabolic interdependence, resource competition, and regulatory responses to environmental conditions.^23, 24^ Among these, cooperative interactions such as reciprocal cross-feeding of metabolites can support collective growth and stability, particularly in consortia composed of auxotrophic strains.^3, 18, 22^ However, the expression and stability of cooperation are highly context-dependent, influenced not only by nutrient availability^44–46^ but also by physiological constraints^32, 33^ and external stressors.^10, 47^ In this work, we showed that antibiotics, typically regarded as cell-targeted stressors, can modulate ecological interactions by altering cellular physiology through the regulation of growth and stress-response pathways.

Previous studies have established that the extent and stability of cooperation in auxotrophic consortia are not fixed but depend critically on environmental conditions, particularly nutrient availability. At low amino acid concentrations, strains depend on each other to meet biosynthetic needs, which makes metabolic exchange essential.^12, 48^ In contrast, when resources are abundant, cooperation can decline or shift to competition or parasitism.^37, 44, 49^ These dynamics are shaped not only by the metabolic costs and benefits of sharing but also by regulatory mechanisms that govern cellular investment strategies under varying nutrient regimes.^50–52^ For example, excess resources can intensify microbial interactions through environmental modification, such as pH changes, reducing community stability,^53^ while spatial structure and local quorum sensing can stabilize cooperative behavior even under changing resource conditions.^54^ Together, these studies highlight the context dependence of cooperation, which is governed by both ecological and physiological constraints.

One of the key regulatory mechanisms underlying this context-dependent cooperation is the stringent response, mediated by ppGpp, which plays a central role in the regulation of growth and metabolism under nutrient limitation or stress. When activated, ppGpp reduces investment in ribosome production while increasing the expression of biosynthetic and stress-related pathways.^31, 55–57^ In auxotrophic communities, this reprogramming can enhance cooperation by promoting the production and export of limiting amino acids.^30, 33, 39, 58^ Thus, the stringent response couples environmental conditions with communal metabolic behavior, shaping cooperation and potentially antibiotic tolerance.^32, 42, 59^

Our results reveal that under amino acid limitation, bacteriostatic antibiotics can paradoxically enhance growth in auxotrophic consortia by reinforcing metabolic cooperation. This effect arises from a physiological feedback loop: inhibition of CHL growth triggers the accumulation of ppGpp, which, in turn, reallocates proteomic resources to amino acid biosynthesis and export. The increased availability of exchanged metabolites promotes cooperation between strains, enabling higher community-level growth compared to untreated controls. This cooperative advantage is specific to nutrient-limited conditions in which metabolic exchange is essential. In these regimes, slow growth becomes beneficial: cells avoid the direct toxicity of translation inhibition while maintaining the exchange of limiting resources. Importantly, the enhanced tolerance observed in co-cultures is not driven by resistance mutations but emerges from the interplay between growth dynamics and metabolic regulation. These findings are consistent with the idea that physiological states associated with slow growth can confer protection against antibiotics that target active cellular processes.^42^ Similar dynamics have been observed in oscillatory cross-feeding systems, where slow-growing populations exhibit enhanced tolerance and cheater suppression.^60^

In conclusion, our study underscores the importance of the ecological context in shaping antibiotic responses of multispecies bacterial communities. The degree of tolerance depends not only on the presence of the antibiotic, but also on nutritional conditions and the composition of the community. Co-cultures respond differently depending on their auxotrophic profiles and the level of amino acid supplementation, suggesting that microbial interactions and physiological regulation jointly shape antibiotic responses. Recognizing that tolerance can emerge from community-level dynamics rather than individual resistance traits broadens our understanding of microbial resilience. Future work should examine how spatial structure, community complexity, and environmental fluctuations influence these dynamics, and whether similar regulatory feedbacks operate under other classes of stressors or in more diverse microbial assemblages. Targeting metabolic interdependencies, rather than individual resistance mechanisms, can offer new strategies to disrupt emerging tolerance and improve antimicrobial efficacy.

## 3 Methods

### Strains and Media

We used *Escherichia coli* auxotrophic strains from the Keio collection,^43^ carrying deletions in Δ*leuB* and Δ*pheA*, rendering them auxotrophic for leucine and phenylalanine, respectively. Each strain was transformed with a plasmid expressing either mCherry or eCFP to allow identification via fluorescence. Synthetic consortia were assembled by combining Δ*pheA* (mCherry) and Δ*leuB* (eCFP). Strains were grown overnight in LB medium supplemented with kanamycin (40 µg/mL) and ampicillin (100 µg/mL) at 37°C. Cultures were then washed with M9 salts (7 g/L K_2_HPO_4_, 2 g/L KH_2_PO_4_, 0.58 g/L Na_3_C_6_H_5_O_7_, 1 g/L (NH_4_)_2_SO_4_, and 0.1 g/L MgSO_4_) to remove residual nutrients, and optical density at 630 nm was adjusted to 0.15–0.18 in M9 minimal medium before inoculation.

To calculate the response to a changing environment we measure the final abundance of the coculture Δ*pheA* (mCherry) and Δ*leuB* (eCFP) in media with leucine and phenylalanine. We grow a 50-50 coculture in M9 glucose supplemented with a range of both amino acids with concentrations 0 to 4 times their basal concentrations increasing 0.25 per condition. All plates were incubated for 48 h at 37°C without shaking. OD and fluorescence were measured at 0, 24, and 48 h using a BioTek Synergy H1 plate reader. The final abundance of each strain was calculated using a Beckman-Coulter flow cytometer with 20,000 events under each environmental condition.

### Dose–Response Experiments

We exposed monocultures and co-cultures to a range of CHL concentrations (0–4 µ*µ*g/mL) in M9 minimal medium supplemented with 2 g/L glucose and varying concentrations of leucine and phenylalanine. Supplementation levels were defined as low (basal), medium (2×), and high (3×) based on predicted amino acid requirements from a genome-scale metabolic model of *E. coli*.^37^ The basal concentrations—16.91 µg/mL for leucine and 8.76 µg/mL for phenylalanine—were calculated using the expression: AC = RDW ·0.295 · ACMW, where RDW is the reference dry weight, 0.295 is the amino acid requirement per unit dry weight, and ACMW is the molecular weight of the amino acid. Cultures were incubated at 37 ^*°*^C with shaking, and OD_630_ and fluorescence were measured at 0 and 48 hours.

### Disk-Diffusion Assay Using Culture Supernatants

To evaluate how environmental conditions and CHL exposure influence the ability of auxotrophic cultures to supply growth-supporting metabolites, we performed a supernatant transfer assay across CHL and amino acid gradients. Auxotrophic *E. coli* strains (Δ*leuB*-eCFP and Δ*pheA*-mCherry) were grown as monocultures or as a 1:1 co-culture in M9 minimal medium supplemented with four amino acid levels (0X, 1X, 2X, 3X) and four CHL concentrations (0, 0.5, 1.5, and 3 *µ*g/mL). Cultures were incubated for 24 h at 37 °C with shaking (250 rpm), with three biological replicates per condition.

After incubation, cells were pelleted by centrifugation (5,000 g, 10 min), and the supernatants were collected and sterile-filtered through 0.22 *µ*m PVDF filters. The growth-supporting capacity of each supernatant was assessed using a disk diffusion assay. M9 agar plates without amino acid supplementation were inoculated either with the co-culture or with the individual auxotrophic strains at OD_600_ = 0.05. Sterile 6 mm paper disks were placed on the agar surface, and 20 *µ*L of each filtered supernatant was applied per disk. Plates were incubated at 37^*°*^C for 24 h, and the radius of the growth zone surrounding each disk was measured using bespoke ImageJ macros.

### Growth Rate Measurements and Fluorescence Quantification

Growth dynamics were evaluated over 48 hours using M9 minimal medium containing either glucose or acetate (2 g/L) and four CHL concentrations (0, 0.5, 1.5, and 3 µg/mL), with all amino acid supplementation levels. OD and fluorescence were recorded every 20 minutes using a BioTek H1 Synergy Hybrid Multi-Mode Microplate Reader. At the end of each experiment, 1 µL of culture was plated on LB agar to assess cell viability. The remaining cells were used to quantify the relative abundance of each strain using an Amnis ImageStream Mk II Imaging Flow Cytometer and a CytoFLEX S Flow Cytometer, analyzing 5,000–20,000 cells per sample. Growth rates were estimated using a Gaussian process-based algorithm implemented in Python.^61^

Specifically, we used Δ*leuB* (leucine auxotroph) and Δ*pheA* (phenylalanine auxotroph), cultured individually or in co-culture in minimal M9 medium with glucose as the sole carbon source. Leucine and phenylalanine were supplemented at defined concentrations based on predicted minimal requirements for monoculture growth, using the iML1515 genome-scale metabolic model of *E. coli*^62, 63^ Basal (low) supplementation levels were set at 16.91 µg/mL for leucine and 8.76 µg/mL for phenylalanine; medium and high levels corresponded to 2× and 3× these basal concentrations, respectively. Increasing amino acid availability alleviated biosynthetic constraints, resulting in higher final cell densities in both monocultures and co-cultures, consistent with faster growth under reduced metabolic limitations. To monitor co-culture composition, we fluorescently labeled each strain with plasmids expressing either mCherry or eCFP, enabling quantification of relative strain abundances by flow cytometry after 48h.

### Dynamic Proteome Partition Mathematical Model

We model proteome allocation, translational inhibition, and amino acid production using a dynamical system that tracks transcription, translation, ppGpp regulation, and CHL inhibition. The state variables are the mRNA concentrations for the metabolic (*m*_*E*_) and ribosomal (*m*_*R*_) sectors; the proteome fractions allocated to metabolic enzymes (*e*_*o*_), amino acid biosynthesis (*e*_*a*_), and active ribosomes (*r*); the concentration of ppGpp (*g*); and the population of inactivated ribosomes (*r*_inh_). The dynamics depend on external concentrations of glucose (*s*), amino acids (*aa*), and chloramphenicol (*CHL*).

The metabolic (*m*_*E*_) and ribosomal (*m*_*R*_) mRNA dynamics depend on nutrient and ppGpp levels:

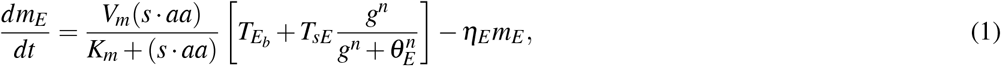

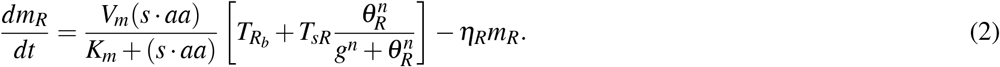

In these equations, *V*_*m*_ and *K*_*m*_ represent the maximum transcription rate and nutrient affinity, respectively. Parameters 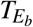 and 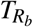 represent basal transcription rates for the metabolic and ribosomal sectors, while *T*_*sE*_ and *T*_*sR*_ denote ppGpp-regulated transcription rates. The Hill parameters *θ*_*E*_, *θ*_*R*_, and *n* define sensitivity to ppGpp, and mRNA degradation occurs at rates *η*_*E*_ and *η*_*R*_.

Translation of mRNAs into proteins leads to changes in proteome composition according to:

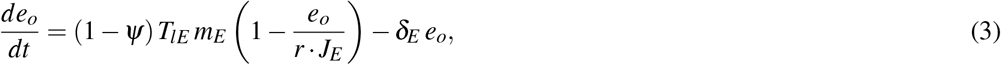

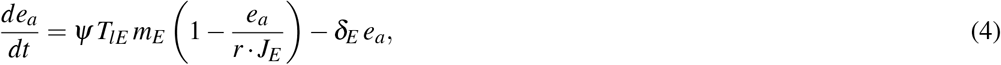

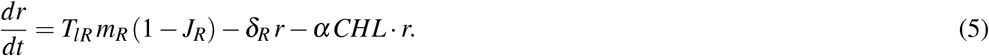

Here, *ψ* denotes the fraction of metabolic translation allocated to amino acid biosynthesis. *T*_*lE*_ and *T*_*lR*_ represent translation rates, and *J*_*E*_ and *J*_*R*_ capture ribosome saturation or translation efficiency. Proteome fractions are degraded or diluted at rates *δ*_*E*_ and *δ*_*R*_, and *CHL* is a bacteriostatic antibiotic that inhibits ribosomes at a rate proportional to both their abundance and the drug concentration.

### Modeling ppGpp Regulation of Resource Allocation

The concentration of the alarmone ppGpp, which regulates proteome partitioning under ribosomal stress, evolves according to:

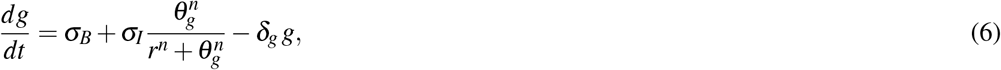

where *σ*_*B*_ and *σ*_*I*_ represent basal and inducible ppGpp synthesis rates, respectively, and *θ*_*g*_ denotes the ribosome threshold for ppGpp induction. The constant *δ*_*g*_ denotes the degradation rate of ppGpp and *n* a Hill coefficient. Thus, ppGpp levels rise when ribosome abundance is low (signaling starvation or inhibition), and decrease when ribosomes are abundant.

To capture the effect of CHL on translation, we further subdivide the ribosomal sector (*φ*_*R*_) into active ribosomes (*r*) and CHL-bound ribosomes (*r*_inh_). Chloramphenicol molecules bind to active ribosomes at a rate proportional to the concentrations of *r* and extracellular antibiotic (*CHL*), forming an inactive complex that is degraded over time:

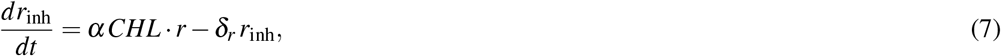

where *α* is the inhibition rate constant and *δ*_*r*_ represents the degradation or dilution rate of inhibited ribosomes.

The total ribosomal proteome fraction is conserved as *φ*_*R*_ = *r* + *r*_inh_. We define *φ*_total_ = *r* + *r*_inh_ + *e*_*o*_ + *e*_*a*_ as the total fraction of the proteome explicitly modeled. The remaining fraction, *φ*_*Q*_, corresponding to growth-independent or housekeeping proteins, is assumed constant and not dynamically tracked.

We assume that amino acid export occurs via passive leakage proportional to biosynthetic flux, so that the effective exchange rate scales with production (the export constant is absorbed into *κ*_*a*_. This matches prior auxotrophic cross-feeding models that treat amino-acid exchange as active uptake coupled to passive leakage from producer cells.^64^ We then define the *growth rate λ* and the *amino acid export rate β* as functions of proteome allocation to the ribosomal and amino acid biosynthesis sectors, respectively:

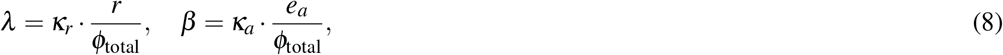

where *κ*_*r*_ and *κ*_*a*_ represent the effective efficiencies of translation and amino acid production. We interpret *λ* as a proxy for cell growth rate and *β* as a proxy for cooperative metabolic activity, specifically amino acid production available for exchange. This normalization ensures that both outputs are expressed per unit of modeled proteome.

### Numerical Experiments

All simulations were performed in Python using the `solve_ivp` function from the `SciPy` library to numerically integrate the system of ordinary differential equations defined in Equations (1)–(7). For each set of environmental conditions, defined by substrate concentration (*s*), amino acid availability (*aa*), and chloramphenicol concentration (*CHL*), the system was integrated over a fixed time interval [0, 1] with 300 time points. Simulations were initialized using the same starting conditions across all runs. The final values of the state variables were used to compute the growth rate *λ* and the amino acid production/export rate *β* as described above. Model parameters are listed in Table S1. The proteome fraction allocated to housekeeping and maintenance functions (*φ*_*Q*_) was assumed to be constant throughout the simulations. All model simulations and figure generation were implemented in a Jupyter notebook, which is available at our GitHub repository: https://github.com/ccg-esb-lab/profileCHL/

## Acknowledgments

We thank D. Romero, A. Escalante, J. Utrilla, and M. Sieber for helpful discussions. We thank S. Ramírez-Barahona for providing the WernerColors library used in this work, and A. Saralegui from the Laboratorio Nacional de Microscopía Avanzada for assistance with flow cytometry. D.R.-G. was a doctoral student in Programa de Doctorado en Ciencias Biomédicas, Universidad Nacional Autónoma de México, and received fellowship 572373 from CONACYT. F.S.-E. received funding from DGAPA-UNAM. H.d.L.-V. was a graduate student in Programa de Maestría en Ciencias Bioquímicas, Universidad Nacional Autónoma de México supported by PAPIIT-UNAM (grant no. IN209419). A.F.-H. was supported by PAPIIT-UNAM (grant no. IN224823). M.R.-G. was supported by the John Templeton Foundation (grant no. 63455) and NSF (grant no. 2316731).

## Supplementary material

**Figure S1.**
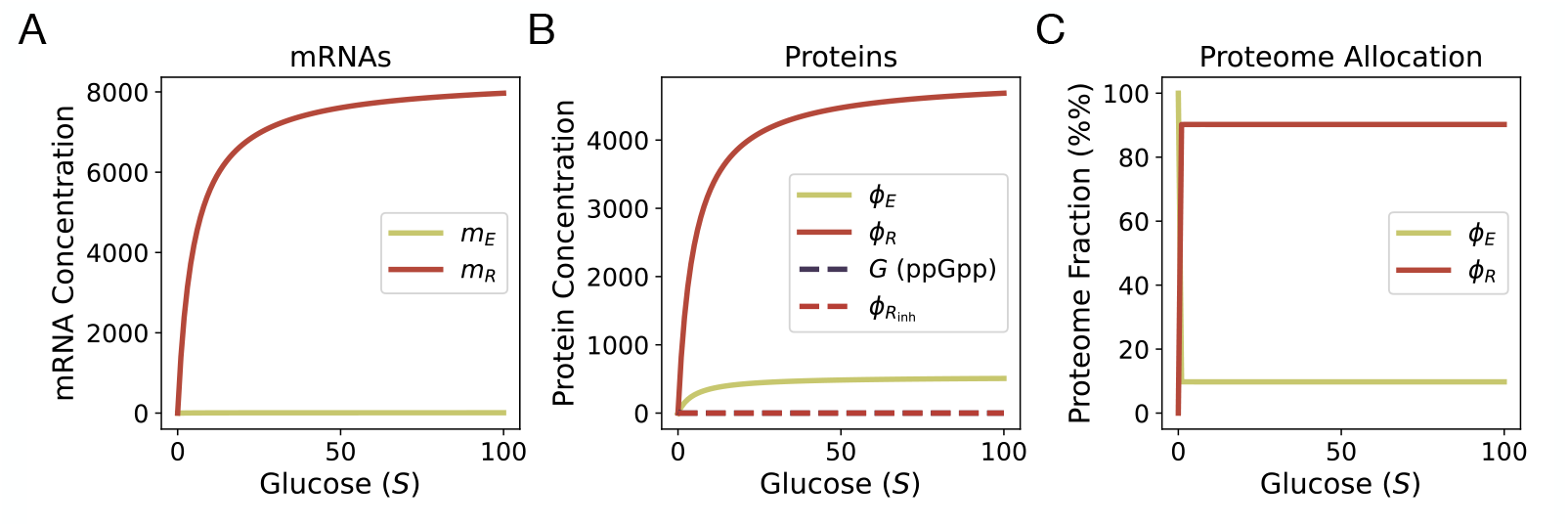
Model response to glucose availability. (A) mRNA concentrations of the metabolic (*m*_*E*_) and ribosomal (*m*_*R*_) sectors as a function of external glucose concentration (*S*). (B) Protein abundances and regulatory factors: metabolic enzymes (*φ*_*E*_, sum of *e* and *aaP*), active ribosomes (*φ*_*R*_), inhibited ribosomes 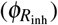, and intracellular ppGpp (*G*). (C) Normalized proteome allocation into metabolic (*φ*_*E*_) and ribosomal (*φ*_*R*_) fractions expressed as percentage of total proteome. Results are obtained from numerical integration of the proteome allocation model across a gradient of glucose supply.

**Figure S2.**
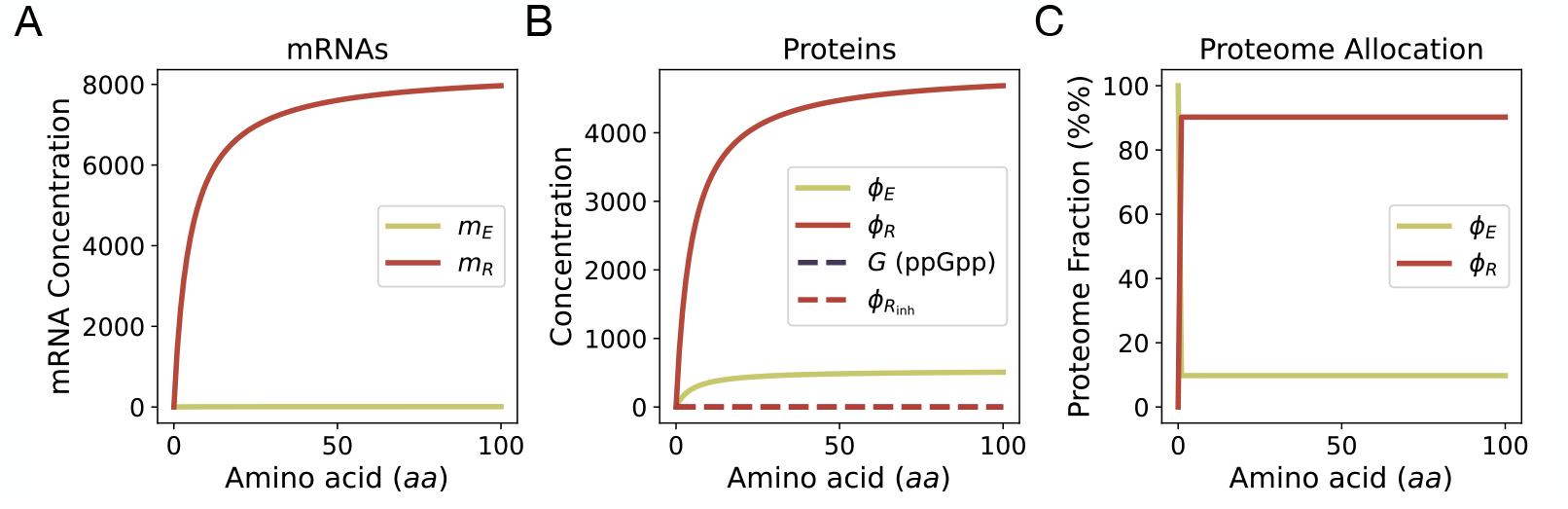
Model response to external amino acid availability. (A) mRNA concentrations of the metabolic (*m*_*E*_) and ribosomal (*m*_*R*_) sectors as a function of external amino acid concentration (*aa*). (B) Protein abundances and regulatory factors: metabolic enzymes (*φ*_*E*_, sum of *e* and *aaP*), active ribosomes (*φ*_*R*_), inhibited ribosomes 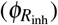, and intracellular ppGpp (*G*). (C) Normalized proteome allocation into metabolic (*φ*_*E*_) and ribosomal (*φ*_*R*_) fractions expressed as percentage of total proteome. Results are obtained from numerical integration of the proteome allocation model across a gradient of amino acid supply.

**Figure S3.**
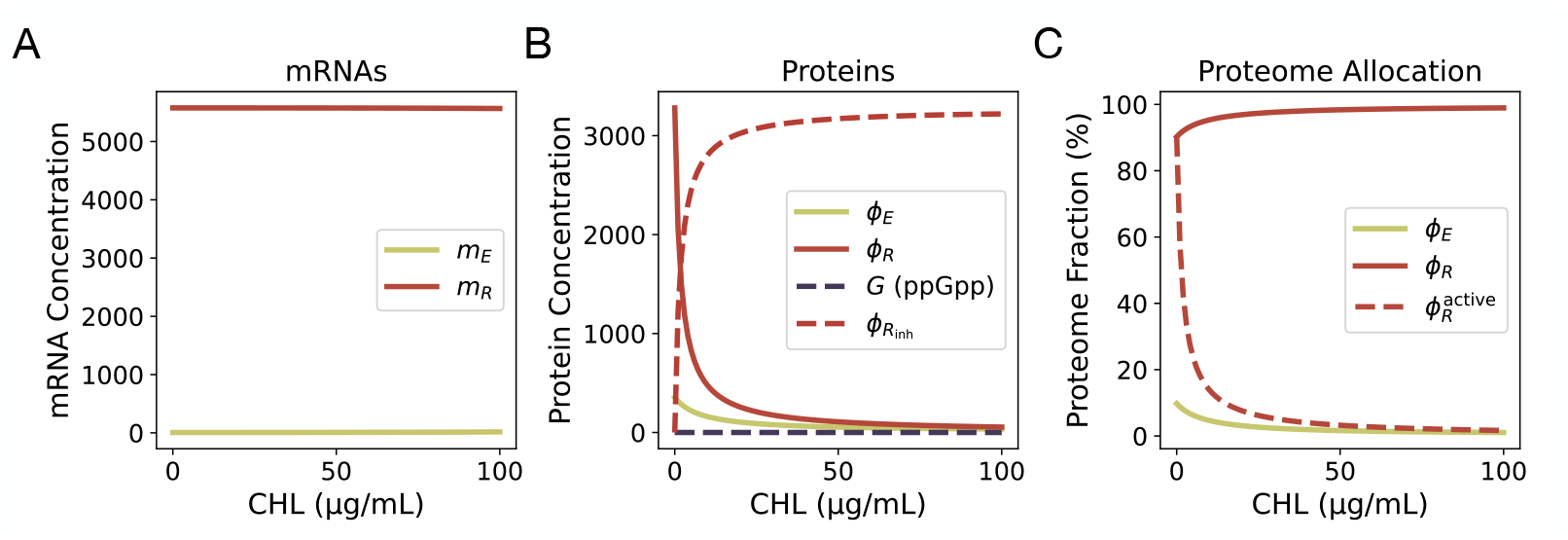
Model response to chloramphenicol exposure. (A) mRNA concentrations of the metabolic (*m*_*E*_) and ribosomal (*m*_*R*_) sectors as a function of CHL concentration. (B) Protein abundances and regulatory factors: metabolic enzymes (*φ*_*E*_, sum of *e*_*o*_ and *e*_*a*_), active ribosomes (*φ*_*R*_), inhibited ribosomes 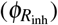, and intracellular ppGpp (*g*). (C) Normalized proteome allocation into metabolic (*φ*_*E*_) and ribosomal (*φ*_*R*_) fractions expressed as percentage of total proteome. Results are obtained from numerical integration of the proteome allocation model across a gradient of CHL concentrations.

**Figure S4.**
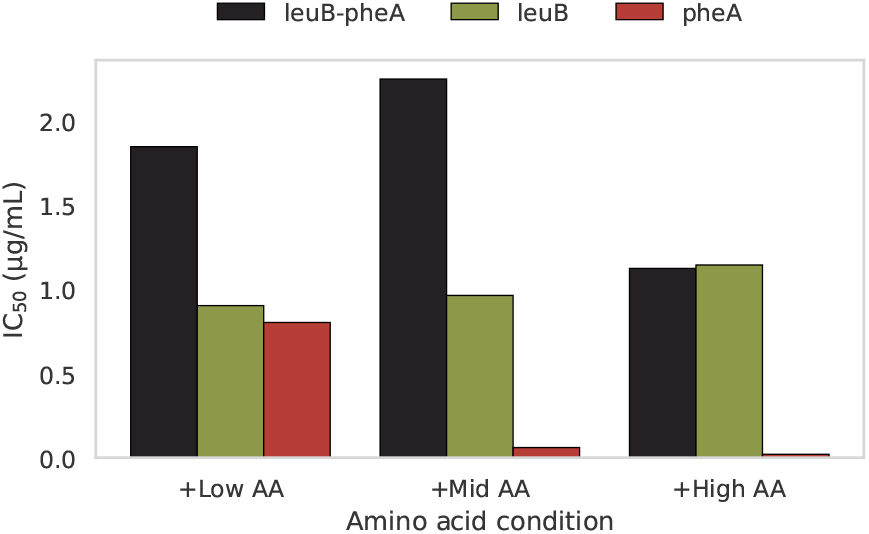
Antibiotic susceptibility measured across amino acid supplementation levels. Bar plots show the chloramphenicol concentration that inhibits final optical density by 50% (IC50) for each culture: the Δ*leuB*–Δ*pheA* co-culture (black) and the corresponding monocultures: Δ*leuB* (green) and Δ*pheA* (red). IC50 values were obtained by fitting dose–response curves across a CHL gradient under three supplementation regimes: Low, Mid, and High.

**Figure S5.**
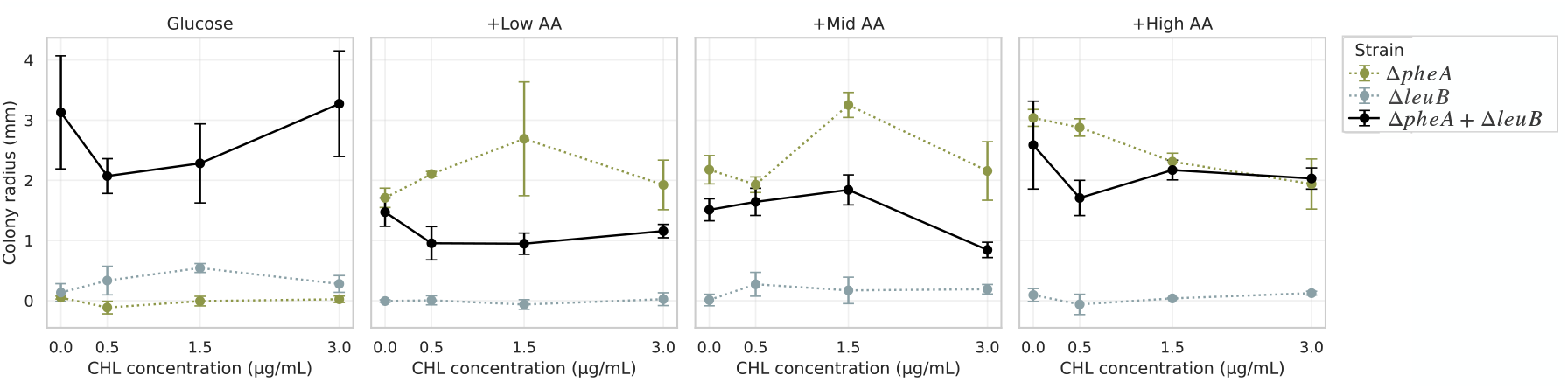
Effect of CHL on the growth-supporting capacity of supernatants across nutrient environments. Colony radius measured on M9 agar without amino acids after applying supernatants collected from 24 h liquid cultures grown across CHL gradients (0, 0.5, 1.5, 3 µg/mL).In amino-acid–supplemented conditions (Low, Mid, High), increasing CHL concentrations led to progressively smaller growth zones for both monocultures, as expected, indicating a dose-dependent reduction in the ability of their supernatants to sustain colony expansion. In contrast, co-culture supernatants generated in unsupplemented glucose displayed a non-monotonic pattern: growth zones were large at 0 µg/mL CHL, decreased at intermediate concentrations, and increased again at 3 µg/mL. This response suggests that strong ribosome inhibition in amino-acid–limited co-cultures increases the concentration of growth-supporting amino acids released into the medium. Points represent mean colony radius; error bars denote SEM across replicates. Subplots correspond to the nutrient conditions used during liquid growth (No supplementation, Low, Mid, High).

**Figure S6.**
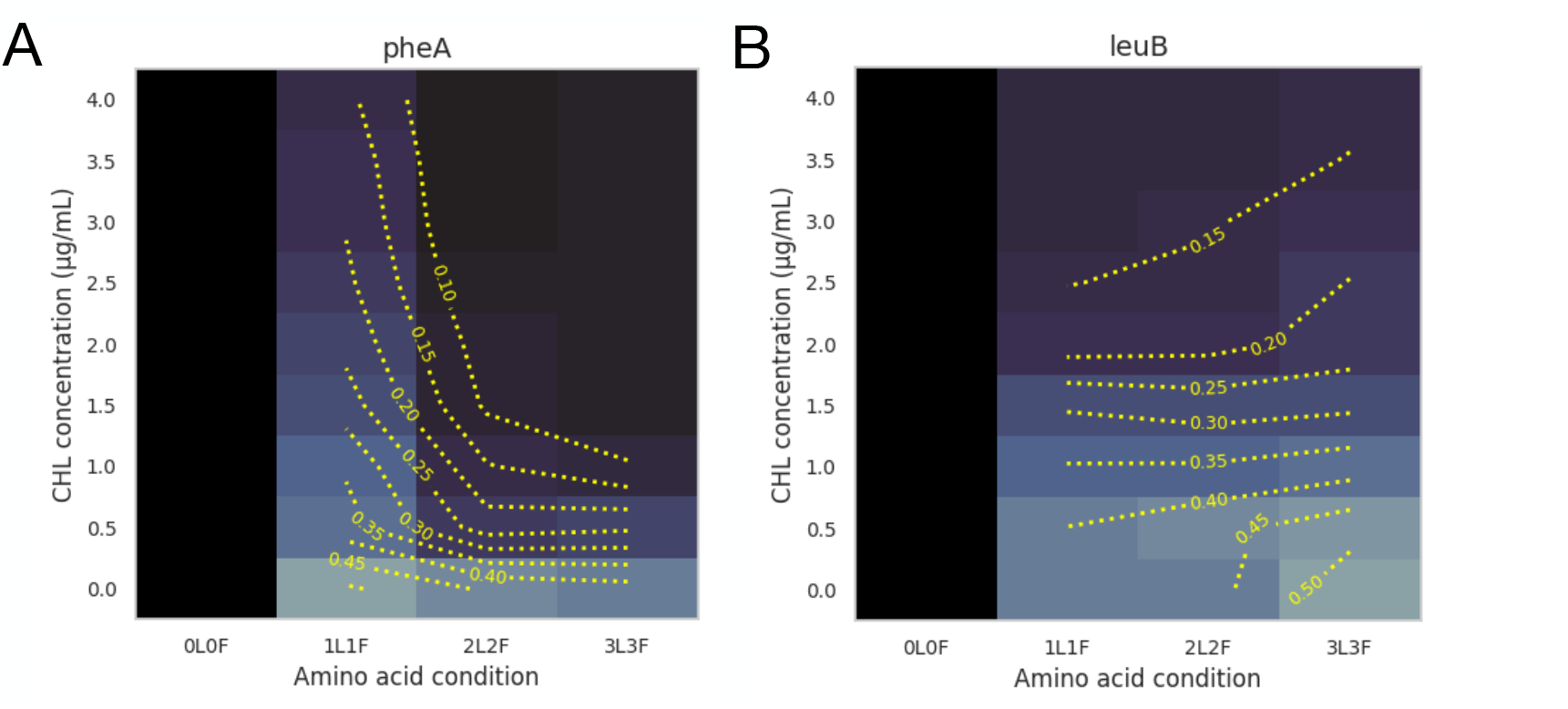
Optical density profiles of auxotrophic monocultures across CHL and amino acid gradients. (A) Optical density (OD) of the Δ*pheA* monoculture after 48 h across a matrix of amino acid supplementation levels (x-axis) and CHL concentrations (y-axis). Heatmaps with yellow contour lines (0.10–0.60) show a uniform, monotonic decrease in OD with increasing CHL under all nutrient conditions. (B) Optical density (OD) of the Δ*leuB* monoculture across the same amino acid and CHL matrix. As in Δ*pheA*, contour lines and heatmap values indicate a consistent monotonic decline in OD with higher CHL, with no growth increase at intermediate concentrations.

**Table S1.** Model parameters used in numerical simulations.

| Symbol | Meaning | Value |
| --- | --- | --- |
| $V_m$ | Maximum transcription rate | $10^3$ |
| $K_m$ | Half-saturation constant for transcription | 50.0 |
| $T_{Eb}$ | Basal transcription rate (E sector) | 0.020 |
| $T_{Rb}$ | Basal transcription rate (R sector) | 0.080 |
| $T_{sE}$ | ppGpp-induced transcription rate (E sector) | 20.0 |
| $T_{sR}$ | ppGpp-induced transcription rate (R sector) | 20.0 |
| $n$ | Hill coefficient | 2.0 |
| $\theta_E$ | ppGpp threshold for E transcription | 5.1 |
| $\theta_R$ | ppGpp threshold for R transcription | 5.1 |
| $\theta_G$ | Ribosome threshold for ppGpp induction | 5.1 |
| $T_{lE}$ | Translation rate of E mRNAs | 80.0 |
| $T_{lR}$ | Translation rate of R mRNAs | 20.0 |
| $\eta_E$ | mRNA degradation rate (E sector) | 2.4 |
| $\eta_R$ | mRNA degradation rate (R sector) | 2.4 |
| $\delta_E$ | Protein degradation rate (E sector) | 1.7 |
| $\delta_R$ | Protein degradation rate (R sector) | 1.7 |
| $\delta_G$ | Degradation rate of ppGpp | 0.8 |
| $J_E$ | Ribosomal saturation constant (E sector) | 0.5 |
| $J_R$ | Ribosomal saturation constant (R sector) | 0.5 |
| $\sigma_B$ | Basal ppGpp synthesis rate | 0.05 |
| $\sigma_I$ | Induced ppGpp synthesis rate | 15.2 |
| $\kappa_r$ | Proportionality constant for growth rate | $1 \times 10^{-11}$ |
| $\kappa_a$ | Proportionality constant for aa production | $0.5 \times 10^{-9}$ |
| $c$ | Resource conversion constant | $13.95 \times 10^9$ |
| $\psi$ | Amino acid proteome cost | 0.01 |
| $\alpha$ | Drug inhibition rate constant | 1.0 |

